# Structural motifs for subtype-specific pH-sensitive gating of vertebrate otopetrin proton channels

**DOI:** 10.1101/2022.03.01.482452

**Authors:** Bochuan Teng, Joshua P. Kaplan, Ziyu Liang, Zachary Kreiger, Yu-Hsiang Tu, Batuujin Burendei, Andrew Ward, Emily R. Liman

## Abstract

Otopetrin (OTOP) channels are proton-selective ion channels conserved among vertebrates and invertebrates and with no structural similarity to other ion channels. There are three vertebrate OTOP channels (OTOP1, OTOP2, and OTOP3), of which one (OTOP1), functions as a sour taste receptor. Whether OTOP channels are gated by, as well as permeating, protons was not known. Here, by comparing functional properties of the three vertebrate proton channels with patch-clamp recording and cytosolic pH microfluorimetry, we provide evidence that each is gated by external protons. OTOP1 and OTOP3 are both activated by extracellular protons, with a sharp threshold of pHe <6.0 and 5.5 respectively, while OTOP2 is negatively gated by protons, and more conductive at alkaline extracellular pH (>pH 9.0). Strikingly, we found that we could change pH-sensitive gating of OTOP2 and OTOP3 channels by swapping extracellular linkers that connect transmembrane domains. Swaps of linkers within the N domain changed the relative conductance at alkaline pH, while swaps within the C domain tended to change the rates of OTOP3 current activation. We conclude that members of the OTOP channel family are proton-gated (acid-sensitive) proton channels and that the gating apparatus is distributed across multiple extracellular regions within both the N and C domains of the channels. In addition to the taste system, OTOP channels are found in the vestibular and digestive systems, where pH sensitivity may be tuned to specific functions.

## Introduction

The activity of ion channels is often tightly controlled through a change in conformation, known as gating, that opens and closes the ion permeation pathway (Hille 2001). Gating can be in response to voltage (voltage-dependent), chemical stimuli (ligand-gated) or membrane deformation (mechanically-gated). Understanding the gating of an ion channel is critical to understanding its physiological function. Recently, a new family of ion channels that are selective for protons and with little or no structural similarity to other ion channels was identified, collectively named the Otopetrins or OTOPs (Tu et al. 2018). These channels mediate proton influx in response to acid stimuli, but whether protons also gate the channels was not known. Importantly, while the structures of vertebrate OTOP1 and OTOP3 channels were recently solved (Saotome et al. 2019; Chen et al. 2019), it is not known if these structures are in open or closed states due to the dearth of information regarding the gating of the channels.

OTOP1 currents were first characterized in taste receptor cells, where pH sensitivity, proton selectivity, and inhibition by Zn^2+^ were described (Bushman, Ye, and Liman 2015; Chang, Waters, and Liman 2010). The founding member of the OTOP family, mOTOP1, was identified as the product of a gene mutated in a murine vestibular disorder (Hughes et al. 2004; Hurle et al. 2003) and was subsequently shown to form a proton channel that functions as a receptor for sour taste in vertebrates (Tu et al. 2018; Zhang et al. 2019; Teng et al. 2019). Most vertebrate genomes encode two related proteins OTOP2 and OTOP3, that also form proton channels (Tu et al. 2018) and are expressed in a diverse array of tissues, including in the digestive tract, where mutations in the corresponding genes have been linked to disease (Tu et al. 2018; Parikh et al. 2019; Qu et al. 2019; Yang, Liu, et al. 2019). Functional OTOP channels are conserved across species, including in invertebrates where they play roles in acid sensing and biomineralization (Hurle et al. 2011; Tu et al. 2018; Mi et al. 2021; Ganguly et al. 2021; Chang et al. 2021).

Ion channels selective for protons are rare in nature, comprising just a small subset of the hundreds of types of ion channels that have been described over the last 80 years (Hille 2001). The two best characterized proton-selective ion channels are M2, a viral protein involved in the acidification of the influenza virus interior, and Hv1, a voltage-gated ion channel that extrudes protons during the phagocyte respiratory burst to maintain pH (Pinto, Holsinger, and Lamb 1992; Ramsey et al. 2006; Sasaki, Takagi, and Okamura 2006; Morgan et al. 2009). OTOP1 channels, unlike HV1, are not gated by voltage (Bushman, Ye, and Liman 2015; Tu et al. 2018) but might instead be gated by protons, like M2 (Liang et al. 2016). However, establishing that its permeant ion gates an ion channel is not trivial. For example, an increase in current magnitudes as pH is lowered could be attributed either to the opening of channels or to an increase in the driving force for proton entry. Here we describe the biophysical response properties of the three murine OTOP channels to varying extracellular pH. By focusing on parameters that are independent of the driving force, we show that extracellular protons gate the three channels in a subtype-specific manner and that the extracellular loop between transmembrane domains eleven and twelve plays a role in gating.

## Results

### Differential current response profiles of three murine OTOP channels to acidic and basic stimuli

We previously reported that the three murine OTOP channels all conduct inward proton currents in response to acid stimuli from pH6-pH4, but with disparate current as a function of pH (I-pH) relations (Tu et al. 2018). We reasoned that because all three channels are permeable to protons, these differences likely reflect differences in gating. To test this hypothesis, we performed a careful comparative analysis of the response properties of murine OTOP1, OTOP2, and OTOP3.

We used patch-clamp recording from HEK-293 cells transfected with cDNA encoding one of the three channels for these experiments. If not otherwise stated, the intracellular solution was pH 7.4 (Cs-Aspartate-based), and the holding potential was -80 mV. All three channels generated inward currents in response to lowering pHo in a Na+-free solution, as previously reported (Tu et al. 2018). The magnitude of the OTOP1 currents increased as the extracellular pH (pH_o_) was lowered over a range of pH 6 to pH 4.5, while the magnitude of OTOP2 currents changed very little over the same pH range (Figure 1A). OTOP3 currents increased more steeply over the same range, with little or no inward currents in response to pH 6 (Figure 1A, C). For all three channels, currents decayed in response to prolonged acid exposure, and the rate of current decay was faster as the pH_o_ was lowered.

**Figure 1.**
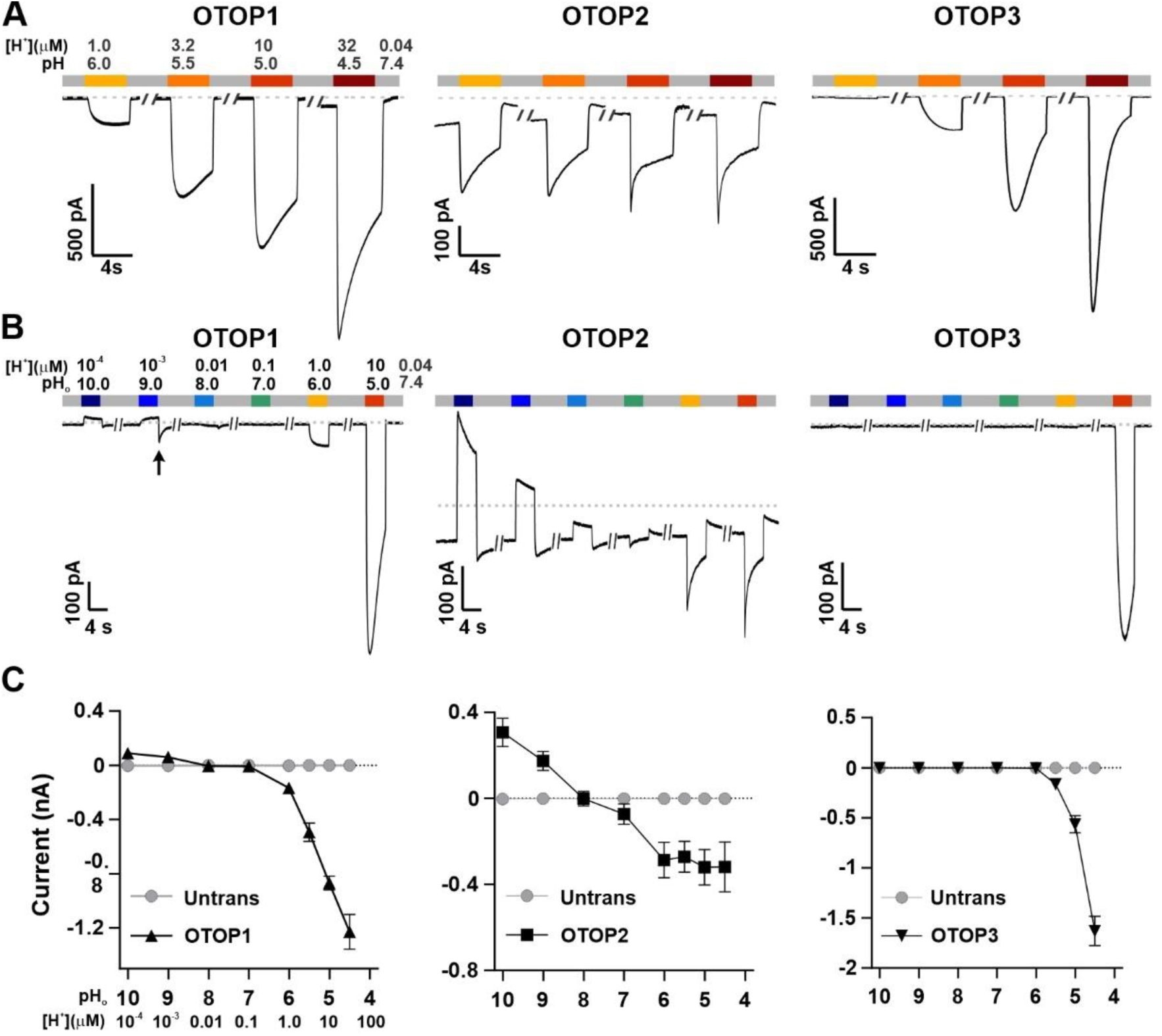
Three vertebrate OTOP channels vary in their response profile across a large pH range. **A**. Proton currents elicited in response to acidic stimuli (pH 6 – 4.5) in the absence of extracellular Na^+^ measured from HEK-293 cells expressing one of the three OTOP channels as labeled (Vm =-80 mV). **B**. Proton currents in response to solutions that varied in pH (pH 10 – 5) measured from HEK-293 cells expressing one of the three OTOP channels as labeled (Vm=-80 mV). **C**. Average data showing the magnitude of proton currents measured from experiments as in (A) and (B). OTOP3 currents are only observed in response to solutions of pH <5.5., while OTOP1 currents are evoked by solutions of < pH 6 and alkaline stimuli, and OTOP2 currents are evoked in response to all stimuli. Data represent mean +/- s.e.m. of biological replicates where for each data point n = 5 – 10 for OTOP1, n = 6 – 10 for OTOP2, and n = 4 – 10 for OTOP3.

To further investigate the response profile of the three channels, we extended the pH range of the test solutions, now including neutral and alkaline solutions (Figure 1B, C). In OTOP1-expressing cells, we observed a small outward current in response to the alkaline solutions (pH 9-10) that was not observed in untransfected cells, and which we, therefore, attribute to currents through OTOP1 channels. In response to the pH 9, but not pH 10 solution, we observed a “tail current” upon return to neutral pH (for pH 9, I_tail_ = -123 +/- 20 pA, n=5; for pH 10, I_tail_ = -36 +/- 11 pA); this suggests that channels had opened in response to the pH 9 stimulus and subsequently closed over a period of several seconds upon return to neutral pH. In response to pH 8 and 7 solutions, no change in the baseline currents was observed.

A very different pH response profile was observed in OTOP2-expressing cells (Figure 1B,C). Quite surprisingly, large outward currents were evoked in response to alkaline solutions of pH 9 and pH 10, which were similar in magnitude to the inward currents evoked in response to the acidic solutions. We also observed changes in the holding current in response to solutions at near-neutral pH (pH 8 or pH 7), suggesting that the channels are open at rest (see below). Overall, we observed that changes in extracellular pH cause changes in OTOP2 currents over the entire pH range tested, and the magnitude of the responses was generally proportional to the driving force on the proton. This relationship broke down over the more acidic pH range, where a change of 10-fold in ion concentration (e.g., pH 6 versus pH 5; Figure 1) did not lead to a substantial increase in current magnitude. Although we do not yet understand this phenomenon, we suspect that it may be due to the rapid decay kinetics of the currents in response to acid stimuli and the consequent attenuation of the peak currents.

OTOP3 currents were active over a narrower pH range than OTOP1 and OTOP2 currents (Figure 1B, C). They were activated steeply below pH 6, and no outward currents were observed in response to any of the alkaline stimuli.

These data suggest that there is a sharp threshold for activation of OTOP1 and OTOP3 channels of ∼pH 6 (OTOP1) and pH 5.5. (OTOP3). In contrast, OTOP2 channels are active over the entire pH range. They also show that both OTOP1 and OTOP2 channels can conduct outward currents, which may be physiologically relevant under some circumstances.

### OTOP2 is selective for protons and open at neutral pH

The unusual response properties of OTOP2 raise the question of whether indeed OTOP2 is selective for protons. OTOP1 is highly selective for H^+^ over Na^+^, by a factor of at least 10^5^ fold (Tu et al. 2018), and currents carried by OTOP3 follow the expectations for a proton-selective current, such as a shift in the reversal potential that follows the equilibrium potential for the H^+^ ion (Tu et al. 2018; Chen et al. 2019). OTOP2 is known to permeate protons (Tu et al. 2018), but selectivity for protons was not previously measured. To address this question, we first performed ion substitution experiments. We observed no change in the magnitude of the currents when the concentrations of either Na^+^ or Cl^-^ in the extracellular solution were changed (Figure 2A), indicating that OTOP2 is not permeable to either ion. We also measured the reversal potential of OTOP2 currents as a function of ΔpH (pHi – pHo) under conditions designed to minimize H+ ion accumulation (see methods) (Tu et al. 2018). These experiments showed that E_rev_ changed ∼59 mV/pH as expected for an H^+^-selective ion channel (Figure 2B). Thus, we conclude that OTOP2 is selective for protons.

**Figure 2.**
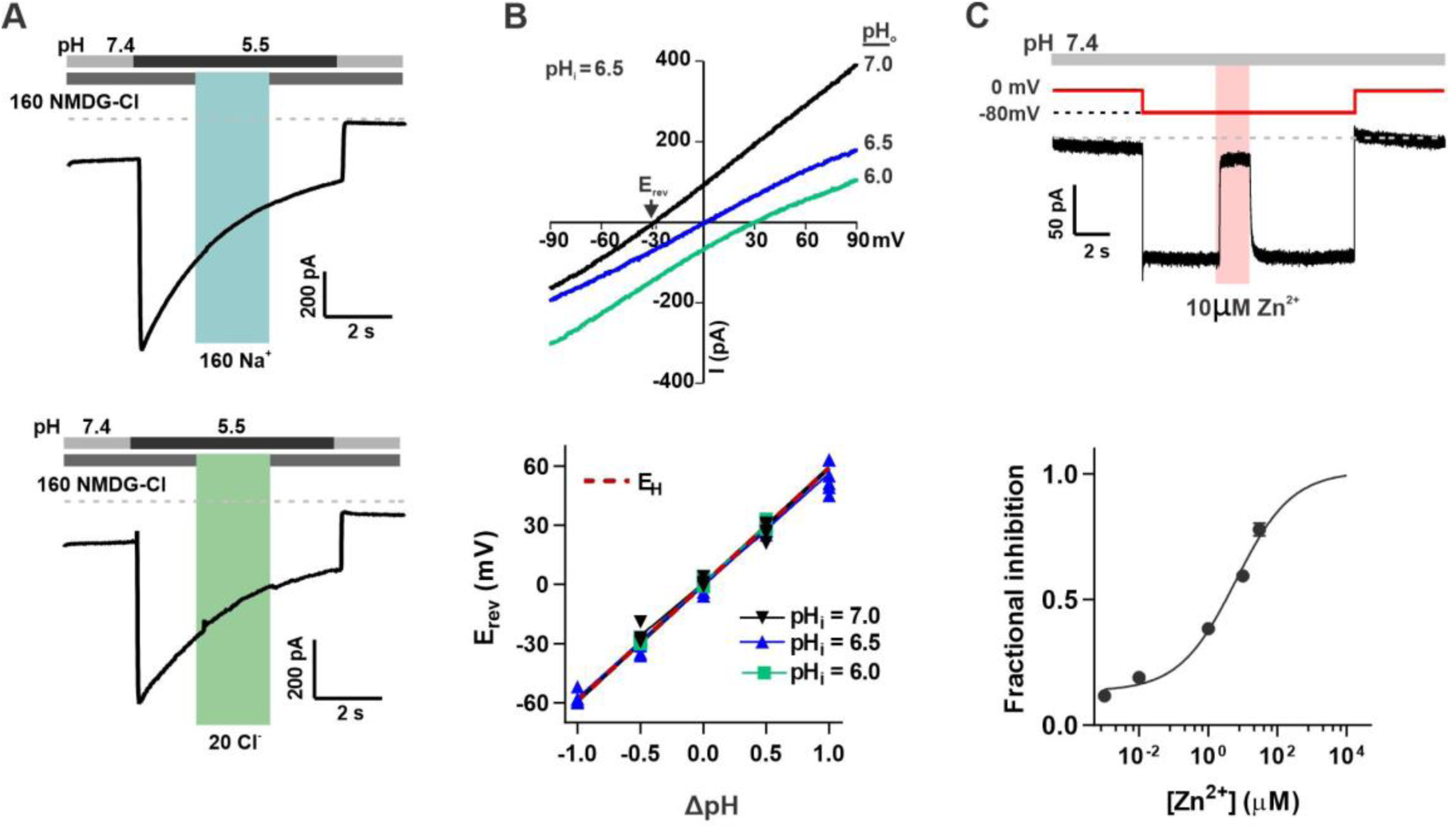
OTOP2 is a proton selective ion channel that is constitutively open at resting pH. **A**. Representative OTOP2 current elicited in response to a pH 5.5 solution with 160 mM Na^+^ replacing NMDG^+^ (top) or 140 mM Methane sulfonate^-^/ 20 mM Cl^-^ replacing 160 mM Cl^-^ (bottom) in the extracellular solutions at times indicated. Vm was held at -80 mV. **B**. The I-V relationship (top) of the Zn^2+^ sensitive component of OTOP2 current in response to different pHo stimuli as indicated obtained from ramp depolarizations in the presence and absence of Zn^2+^. pHi was adjusted to 6.5. Bottom: Erev measured in as a function of ΔpH (pHi – pHo). pHi was adjusted to 6.0, 6.5, or 7.0 as indicated. The red dotted line is the predicted equilibrium potential for H+, EH. **C**. Extracellular Zn^2+^ inhibits resting OTOP2 currents in a dose-dependent manner. Trace (top) shows inhibition of resting current in OTOP2-expressing HEK-293 cells by 10 µM Zn^2+^. Vm as indicated. Fractional block was fit with a Hill slope = 0.53 and IC50 = 6.1 µM. Data represent mean +/- s.e.m of biological replicates where n = 3 – 4 for each data point.

The unusual response properties of OTOP2 suggested that it might be open at rest. For example, we routinely observed a large resting current at -80 mV in OTOP2-expressing cells. We reasoned that if this resting current is due to open OTOP2 channels, it should be inhibited by Zn^2+^, a blocker of OTOP1 and other proton channels (Tu et al. 2018; Decoursey 2003). Indeed, OTOP2 currents at neutral pH were inhibited by Zn^2+^ with an IC_50_ of 5.6 µM (Fig 2C). As the inhibition of OTOP1 by Zn^2+^ is pH-dependent (Teng et al. 2019; Bushman, Ye, and Liman 2015), the relatively high-affinity block of OTOP2 is not unexpected.

Thus, we concluded that OTOP2 is a proton-selective ion channel that is open at neutral pH.

### Different pH-response profiles of three murine OTOP channels measured with pH imaging

Inward and outward currents carried by protons through OTOP channels are expected to cause a change in intracellular pH. Thus, to confirm the results described above, we assessed the activity of each OTOP channel by monitoring intracellular pH. For these experiments, we co-transfected HEK-293 cells with a pH-sensitive fluorescent protein – pHluorin, a variant of GFP whose fluorescence emission changes as a function of the intracellular pH (Miesenbock, De Angelis, and Rothman 1998). As a control to confirm the expression and activity of pHluorin, all cells were exposed at the end of the experiment to acetic acid, which crosses cell membranes and causes intracellular acidification (Figure 3A -- D, dark red bar) (Wang et al. 2011).

**Figure 3.**
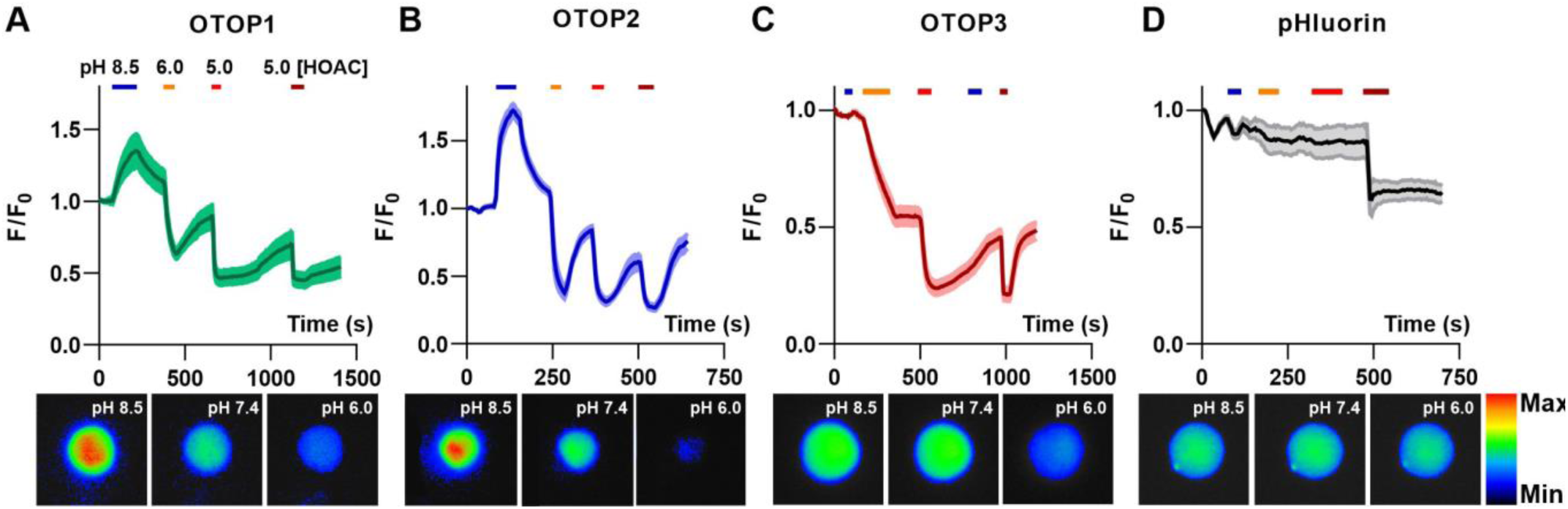
OTOP channels mediate influx and efflux of protons as measured with fluorescence emission from pHluorin. **A-D**. Changes in fluorescent emission upon exposure to varying extracellular pH, as indicated, measured from HEK-293 cells co-expressing OTOP channels and the pH-sensitive indicator pHluorin (A-C) or pHluorin alone (D). Data are shown as the mean ± SEM for n = 7, 7, 11, and 3 cells for (A), (B), (C), (D) respectively. Acetic acid, which is permeable through cell membranes and acidifies cell cytosol directly, served as a positive control. Only OTOP1 and OTOP2 conducted protons out of the cell cytosol in response to alkalinization, while all three channels conducted protons into the cell cytosol in response to acidification, albeit at different rates. The lower panel shows images of a single cell in the field of view used for these experiments taken at the pH indicated (pseudo color is arbitrary units).

Cells expressing each of the three OTOP channels responded to the acidic solutions with large changes in intracellular pH, a response not observed in control cells (Figure 3A – D). Strikingly, we also observed changes in intracellular pH in response to the alkaline extracellular solution (pHo = 8.5) in both OTOP1- and OTOP2-expressing cells, but not in OTOP3-expressing cells. This data is consistent with the patch-clamp data showing that OTOP1 and OTOP2 channels conduct outward proton currents in response to alkaline extracellular stimuli, while OTOP3 channels do not conduct outward currents under those conditions.

We also noted differences in the time course of the response to acidic solutions (pH 6 or pH 5) in cells expressing the three OTOP channels. Notably, the response of OTOP3-expressing cells was significantly slower than that of OTOP1 or OTOP2-expressing cells. This is likely a consequence of the smaller currents that are evoked by these stimuli in OTOP3 cells and the slower kinetics of channel activation (see below).

Together these data support the conclusion that OTOP1 and OTOP2 can conduct outward proton currents in response to alkaline extracellular solution solutions, while all three channels carry inward proton currents in response to acidic stimuli.

### Kinetics of OTOP channels provide direct evidence of pH-dependent gating

In patch-clamp recordings such as those shown in Figure 1, we noted differences between the three OTOP channels in the kinetics of the currents elicited in response to acidic solutions. For example, OTOP2 currents showed faster activation than OTOP1 currents, while OTOP3 currents were considerably slower. As the solution exchange was the same for all three channels, differences in the rate of activation of the currents can only be explained by differences in the gating of the channels. In particular, for a channel that is constitutively open (OTOP2), we expect the currents to change in response to the change in concentration of the permeant ion (H+) with a time course that reflects the speed of the solution exchange. In contrast, for a channel closed at neutral pH (OTOP1 and OTOP3), currents may increase more slowly, with kinetics that are dependent on the opening rate of the channels (Figure 4 A).

**Figure 4.**
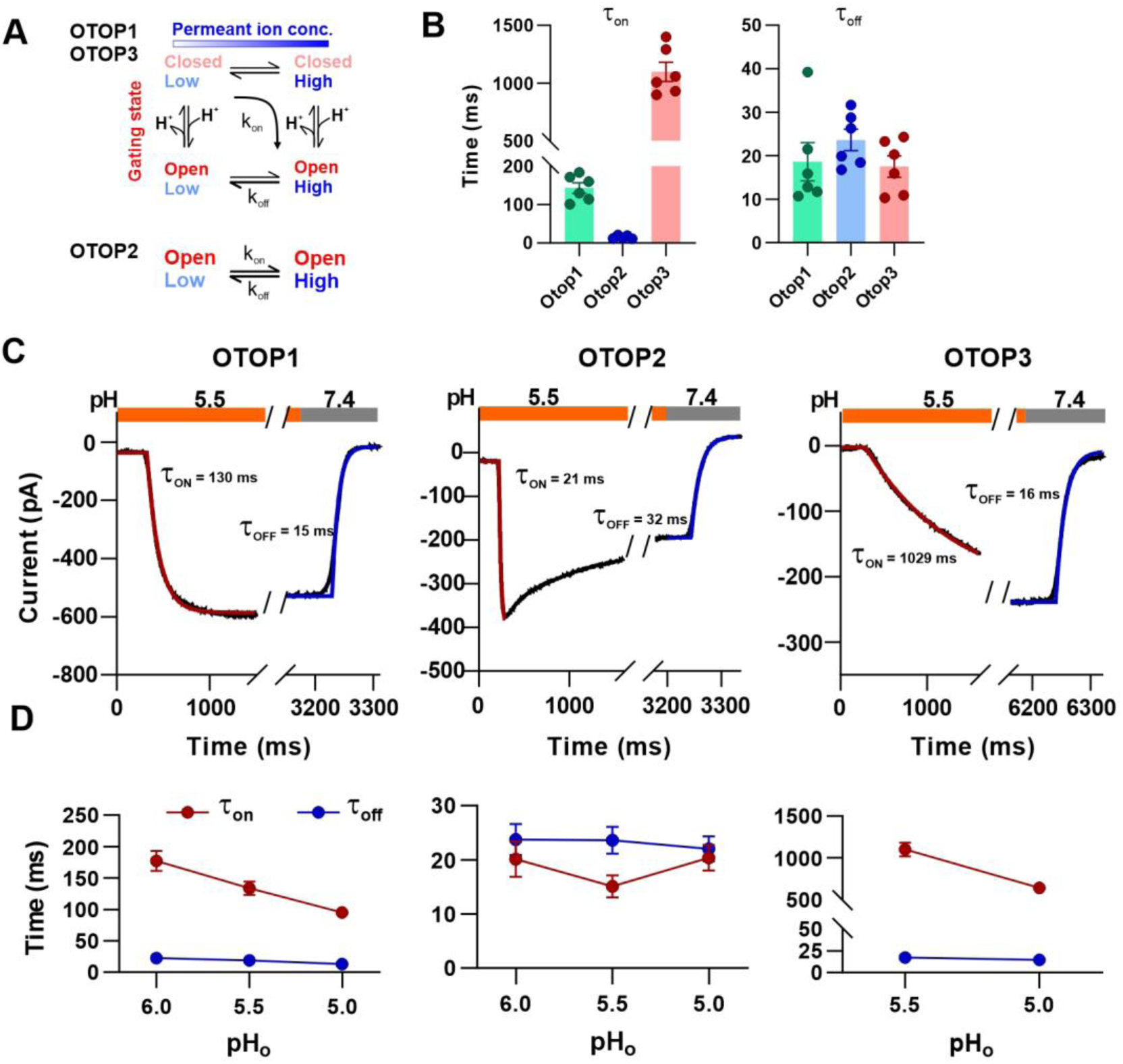
Activation kinetics vary dramatically between OTOP channels. **A**. Simplified model showing how gating and changes in the concentration of permeant ions combine to determine the kinetics of currents carried by OTOP channels in response to changes in extracellular protons. OTOP1 and OTOP3 channels must undergo a pH-dependent gating transition, which if slow enough, will dominate the kinetics of the on rate. OTOP2 channels do not need to undergo a gating transition and on rate for currents is determined solely by the rate of the solution exchange. For all three channels, off rates are determined by the rate of solution exchange (transition from high to low concentration of permeant ions). **B**. Summary data for τon (left panel) and toff (right panel) of the three OTOP channels (OTOP1: green, OTOP2: blue, OTOP3: red) in response to the application and removal of pH 5.5 extracellular solution. n = 6, 5-6, 6 for OTOP1, OTOP2 and OTOP3 respectively (τon and toff). **C**. Representative current traces (black) from cells expressing each of the three OTOP channels in response to the application and the removal of pH 5.5 solutions. Acid stimuli were applied at time = 0; the lag in the activation and deactivation represents the time for solution exchange. The activation and the decay of the currents were fitted with single exponential curves (red and blue curves, respectively). **D**. Time constants of the activation (τon, red) and the deactivation (τoff, blue) of OTOP1, 2 and 3 currents in response to acidic stimuli (pH 6.0, 5.5 and 5.0) measured from experiments as in C. Note that the data at pH 5.5. is also shown in the scatterplot in Panel B. n = 6,5-6, and 6 for OTOP1, OTOP2 and OTOP3, respectively (τon and toff).

The rate of activation of each channel in response to acid stimuli was measured by fitting the time course of the current upon solution exchange with a single exponential. Note that we excluded the first few milliseconds where responses deviated from an exponential time course, likely due to the non-instantaneous rate of solution exchange. We also measured the rate of decay of the currents after return to neutral pH in a similar manner.

In response to the pH 5.5 stimulus, the activation kinetics of the three OTOP channels varied across nearly two orders of magnitude (Figure 4B, C). The time constant for activation of OTOP2 was 15.1 ± 2.0 ms (n = 6) (Figure 4B, C), while time constants for activation of OTOP1 and OTOP3 were 142.7 ± 13.5 ms (n = 6) and 1140.1 ± 81.7 ms (n = 7) (Figure 4B, C), respectively. The observation that currents carried by OTOP1 and OTOP3 are activated with a time course that is considerably slower than the solution exchange suggests that extracellular protons gate them. In contrast, and as expected, the time constant for deactivation of OTOP1, OTOP2, and OTOP3 currents were very similar (τ = 18.6 ± 4.4 ms, n = 6, 20.5 ± 3.7 ms, n = 7, and 17.0 ± 2.2 ms, n = 7, respectively) (Figure 4B, C), reflecting the rate of solution exchange and removal of the permeant ion (Figure 4A). This also confirms that the differences in on-rates were not artifacts of varying solution exchange times between groups of cells.

We also measured the kinetics of the currents carried by all three channels in response to solutions that varied in pH. The activation rate of OTOP1 and OTOP3 currents increased as the pH was lowered. This is as expected if protons gate the channels. In contrast, the activation rate of the OTOP2 currents was insensitive to the extracellular pH over this range, consistent with an interpretation that lowering the pH does not open the channels, which are already open (Figure 4D and Figure S1). In contrast, the deactivation rate of the currents upon return to neutral pH did not vary as a function of pH for any of the three channels and instead reflected the exchange time for the solutions, as expected (Figure 4D).

Together, the slow, pH-dependent kinetics of currents carried by OTOP1 and OTOP3 channels provides strong evidence that they are gated by extracellular protons.

### Effect of extracellular pH on slope conductance

To provide further evidence that extracellular protons gate the OTOP channels, we measured conductance (I/V) as a function of pH for all three channels. Note that this provides a measure independent of the driving force for proton entry, which necessarily changed during these experiments. The macroscopic conductance that we measure is related to channel open probability (Po) by the equation G= N*Po*g, where N is the number of channels and g is the single-channel conductance. Since the number of channels is constant, the slope conductance is a measure of Po*g. We recognize that g may change under different ionic conditions. The slope conductance was measured from -80 mV to 0 mV using ramp depolarization applied once per second. Because the currents decayed during the stimulus, we used the slope of the first ramp at or shortly following the maximum response to each stimulus (pHo = 10 to 5) (Figure 5 A, B).

**Figure 5.**
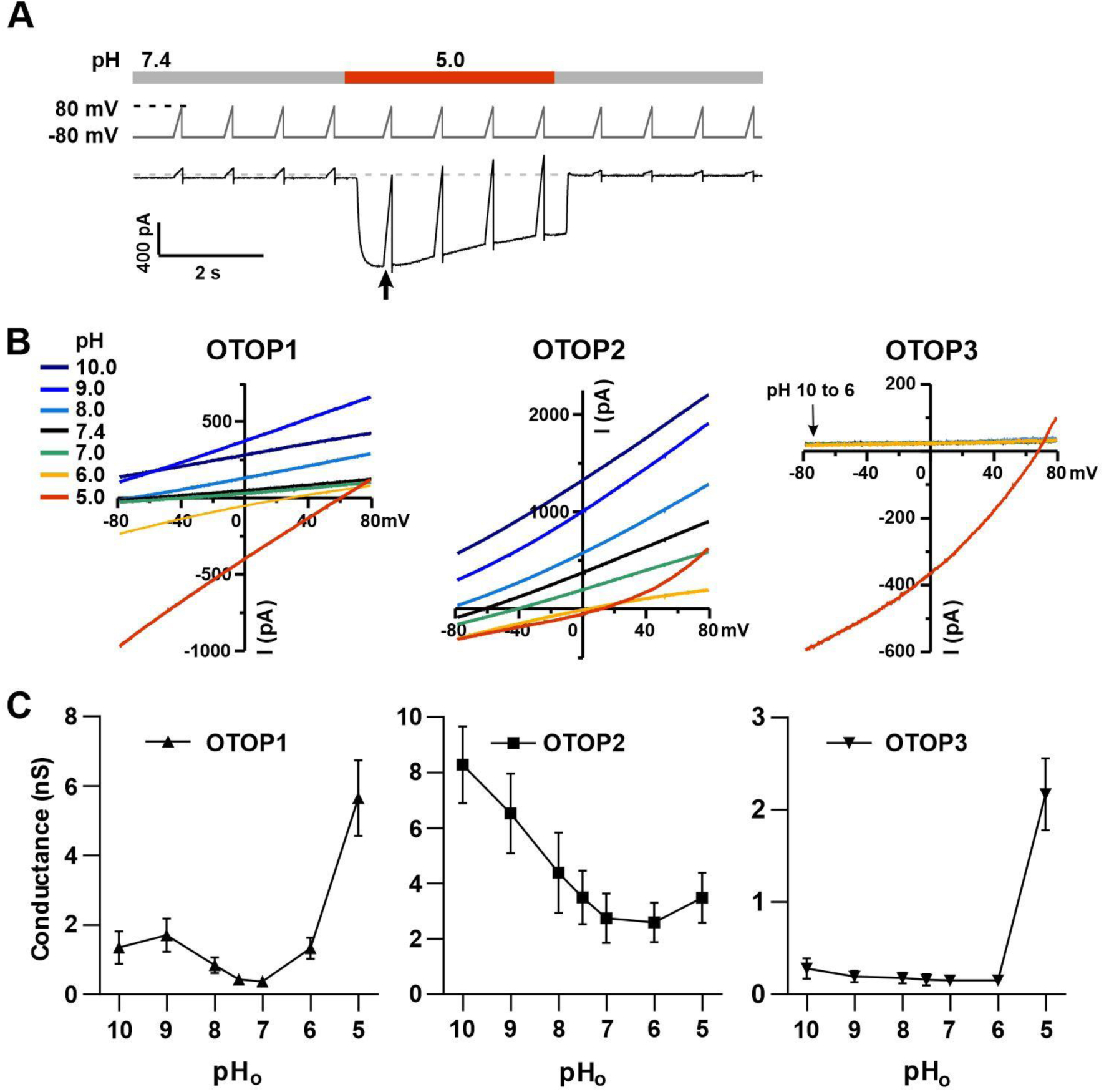
Changes in the slope conductance of OTOP channels as a function of extracellular pH. **A**. Voltage and solution exchange protocol designed to measure the slope conductance over a range of extracellular pHs. Vm was held at -80 mV and ramped to +80 mV (1 V/s at 1 Hz). The first ramp after the currents peaked was used for later measurements. **B**. Representative current-voltage relationship from HEK-293 cells expressing each of the three OTOP channels in response to alkaline or acidic stimuli (pH 10 to 5) from experiments described in (A). The conductance was measured from the slope of the I-V curve between - 80 mV and 0 mV to avoid contamination from outwardly rectifying Cl^-^ currents. **C**. Average slope conductance measured from cells expressing each of the three OTOP channels in response to different pH stimuli from data as in (B).

In OTOP1-expressing cells, the slope conductance varied as a function of extracellular pH, with a dramatic increase in the slope conductance when the extracellular pH was lowered below pHo = 6.0 (Figure 5 B, C). We also observed an increase in the slope conductance when the pH was made more alkaline, consistent with data in Fig 1 showing that an alkaline pH of 9.0 can be activating. In OTOP2-expressing cells, the slope conductance was highest in alkaline pH (pHo = 10) and decreased as the pHo was lowered to pHo = 6, and then slightly increased when pHo was further lowered to 5 (Figure 5 B, C). The response of OTOP3-expressing cells was more similar to that of OTOP1, but with some clear differences (Figure 5 B, C). The slope conductance remained very small when extracellular pH was between 10 and 6 and increased drastically when pHo was 5.

### Extracellular loops are key determinants for pH-sensitive gating

The difference in pH sensitivity of the three murine OTOP channels provided an opportunity to begin to identify elements of the gating apparatus. Specifically, we generated chimeras between mOTOP2 and mOTOP3, which as described above, are the most divergent functionally of the three murine OTOP channels. OTOP channels contain twelve transmembrane domains (S1-S12), with N and C termini located intracellularly. The transmembrane domains 1-6 and 7-12 respectively constitute the structurally homologous “N” and “C” domains. Reasoning that the residues involved in sensing the extracellular pH and gating the channels would be located on the extracellular surface of the channels, we swapped each of the six extracellular linkers that connect transmembrane helices. Each of the twelve chimeric channels was then tested over a range from pH 10 to pH 5. To simplify the analysis, we divided the chimeras into four categories: OTOP2 N domain (OTOP2 backbone with OTOP3 linkers), OTOP2 C domain, OTOP3 N domain, and OTOP3 C domain.

Of the three OTOP2 N domain chimeras, two were functional. Strikingly, the introduction of the OTOP3 S5-S6 linker (L5-6) nearly eliminated the outward currents in response to alkaline stimuli, and reduced the response to the mildly acidic stimulus (pH 6), but had little effect on the magnitude or kinetics of the inward currents elicited in response to the pH 5 stimulus (Figure 6 A-C; OTOP2/OTOP3(L5-6). Introduction of the OTOP3 S1-2 linker reduced current magnitudes (possibly due to effects on trafficking) but did not significantly change relative responses to the stimuli of varying pH. In contrast to the OTOP2 N domain chimeras, the two functional C domain chimeras (L7-8 and L11-12) showed a similar pH-dependence as WT OTOP channels, suggesting that they did not specifically affect channel gating (Figure 6 A, D-E). None of the chimeras showed a significant change in kinetics (Figure 7 A, B, E).

**Figure 6.**
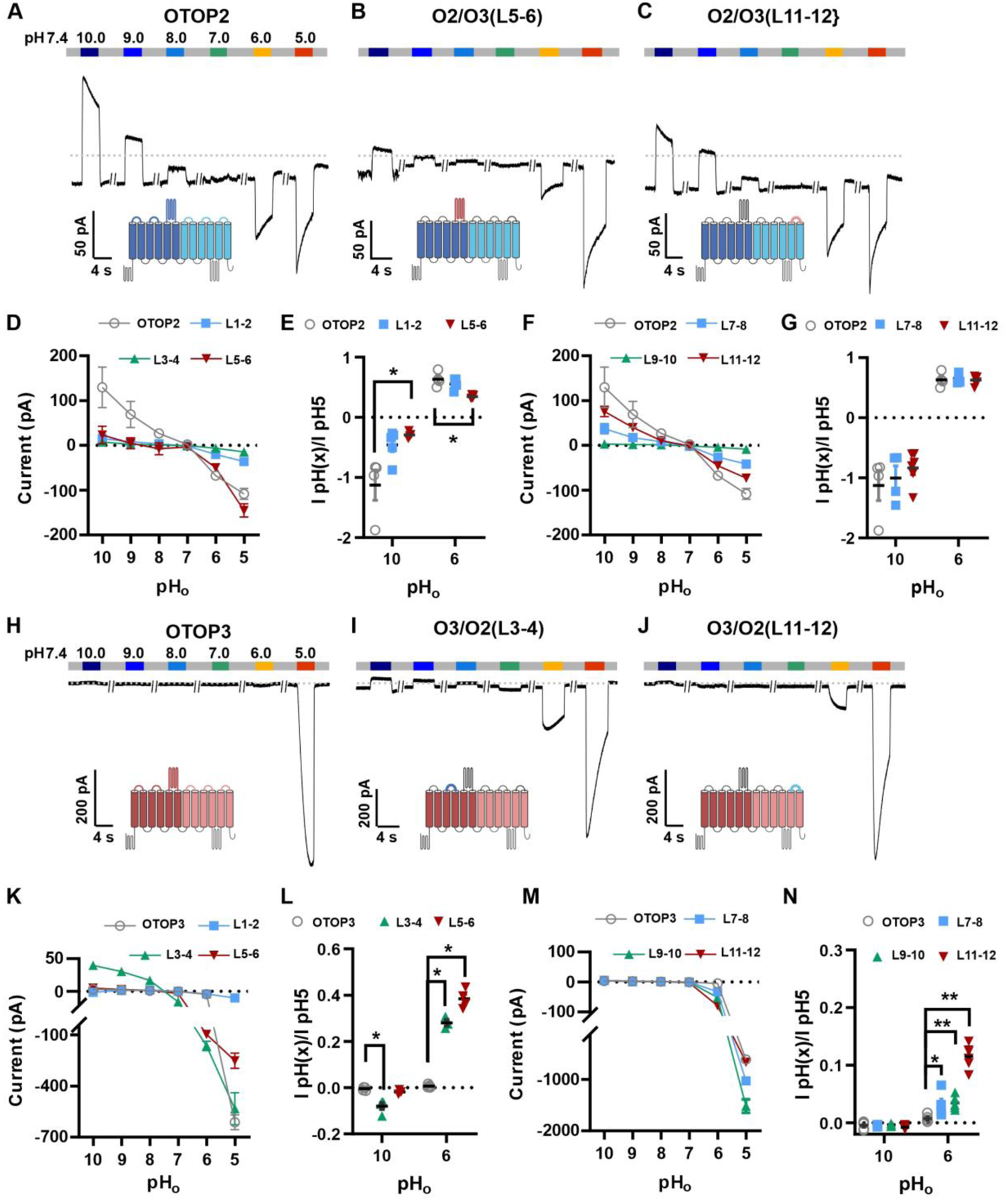
Chimeric channels with external linkers swapped reveal potential gating modules. **A, F**. Representative current traces in response to varying pH from 10-5 measured from HEK-293 cells expressing OTOP2 channels and chimeras with an OTOP2 backbone (**A**) or OTOP3 channels and chimeras with an OTOP3 backbone (F) where numbers refer to linkers between corresponding transmembrane domains. Membrane potential was held at -80 mV. **B, D, G, I**, Current magnitude (mean +/- s.e.m.) for OTOP2 N-domain chimeras (n=4-5) (B), OTOP2 C-domain chimeras (n=4-10) (D), OTOP3 N-domain chimeras (n=3-5) (G), and OTOP3 C-domain chimeras (n=4-6) (I) from experiments such as in (A) and (F). **C, E, H, J**. Same data for pH 10 and pH 6, normalized to the response to pH 5.0 to control for differences in expression. Significance[2w4a]s tested using the Mann-Whitney test. The P-values and n are given in supplementary Table.1.

**Figure 7.**
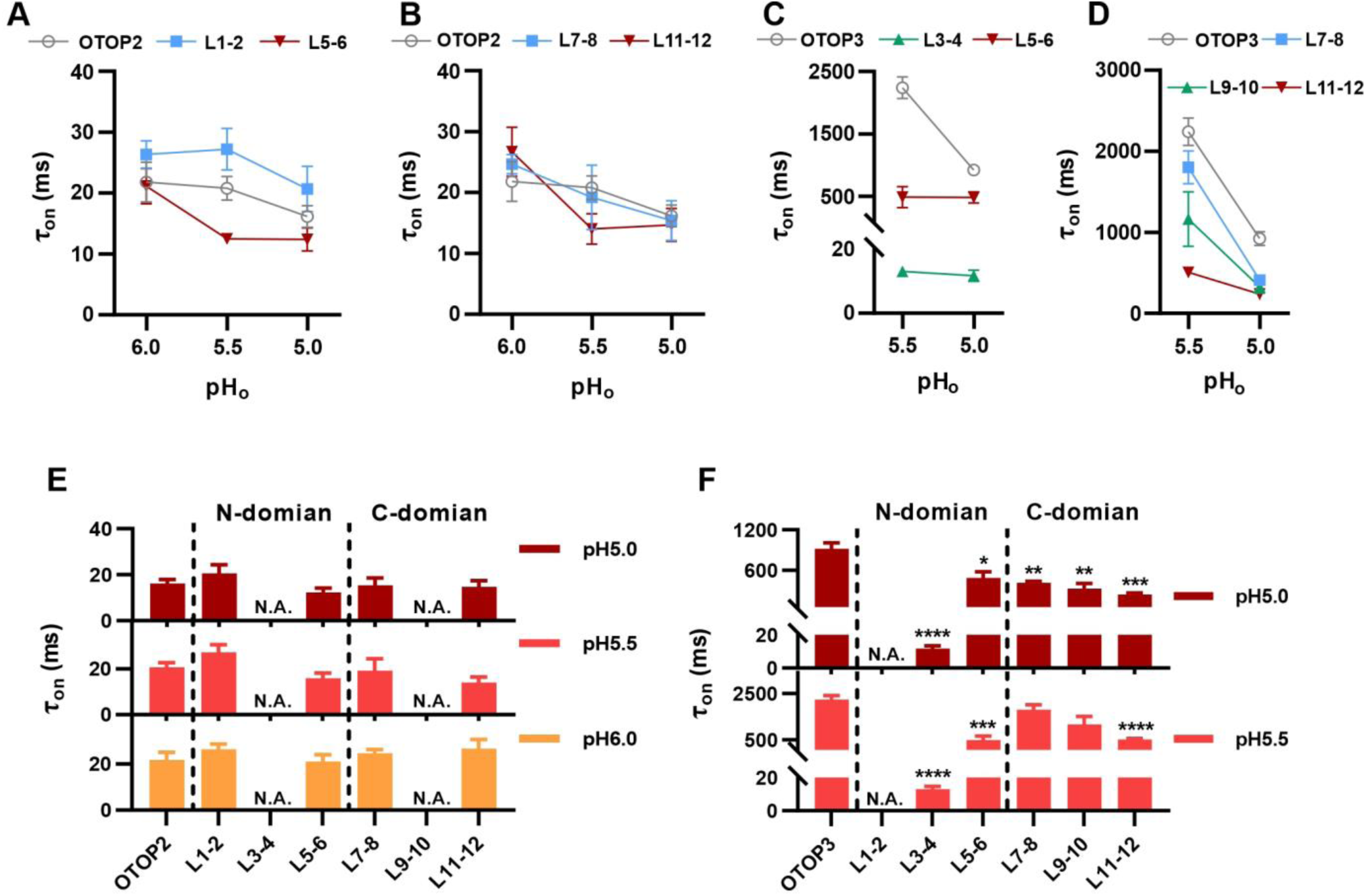
Increase in activation kinetics of mOTOP3 chimeric channels. **A-D**, Time constants for activation of chimeric as compared with wildtype channels, measured from traces as in Fig 6 A,F, using methods as in Fig 4. The τ_on_ of OTOP2 and its chimeras (A, B) was not pH dependent and followed the rate of the solution exchange. Data are mean +/- s.e.m, (n=3-4). The τ_on_ of OTOP3 chimeras bearing OTOP2 linkers(C,D) was generally faster than that of the wildtype channels (n=3-4). Same nomenclature as in Fig 6. **E**,**F**. Same data as in A-D, plotted to allow comparison across all chimeras at the same pH (as indicated; note that because wildtype OTOP3 is not activated by pH 6, data using a pH 6 stimulus is not included in the analysis). Statistical analysis comparing chimeric channels with wildtype channel with Student’s T-test. *, p<0.05, **, P<0.01, ***, p<0.001, ****, p<0.0001

Next, we examined OTOP3 channels with extracellular linkers from OTOP2. Of these, the most striking was an OTOP3 N domain chimera with the L3-4 of OTOP2 which conducted outward currents in response to extracellular alkalinization (Figure 6 F-H). Moreover, this chimera was activated at a more mild pH than OTOP3 channels and with faster kinetics of activation (Figure 7 C, F). Thus, the simple swap of the 3-4 linker conferred a partial gating phenotype like that of the donor, OTOP2. The chimera containing the L5-6 from OTOP2 was more similar to OTOP3, while the L1-2 chimera appeared to be non-functional. Interestingly, the C domain chimeras also produced apparent changes in gating (Figure 7D, F): all three chimeras activated more rapidly and at higher pH, adopting some features from the donor channel (OTOP2). But they retained the steep pH sensitivity of the OTOP3 channels and did not support proton efflux in response to alkaline stimuli like OTOP2.

The structure of OTOP2 channels has not yet been resolved experimentally, and structures OTOP1 and OTOP3 channels do not allow the visualization of the S5-6 linker (Saotome et al. 2019; Chen et al. 2019), possibly because it adopts multiple conformations. Thus, to gain structural insight into the possible mechanisms by which extracellular linker regions could contribute to gating, we examined structures of OTOP channels predicted by Alphafold (Varadi et al. 2022; Jumper et al. 2021). Inspection of the predicted structures of the three mammalian OTOP channels shows the putative location of the extracellular linker and associated transmembrane domains (Figure 8). Interestingly, the 5-6 linker is longer and transmembrane domain 5 (TM5) of OTOP2 adopts a helical structure with a pronounced kink, where that of OTOP3 is straight and rod-like. Differences in the 3-4 linkers and associated transmembrane domains are evident but less pronounced. Thus, it is possible that the introduction of the mOTOP3 S5-6 linker onto mOTOP2, which disrupts conduction at alkaline pH, may do so through a structural rearrangement of transmembrane helices within the N domain of the channel. In contrast, the C terminal linkers are more similar between mOTOP2 and mOTOP3, suggesting that single amino acids that differ between the channels may tune the pH dependence of gating. Together, we conclude that extracellular linkers participate in the gating of OTOP channels and that the gating apparatus is distributed across multiple extracellular regions within both the N and C domains.

**Figure 8.**
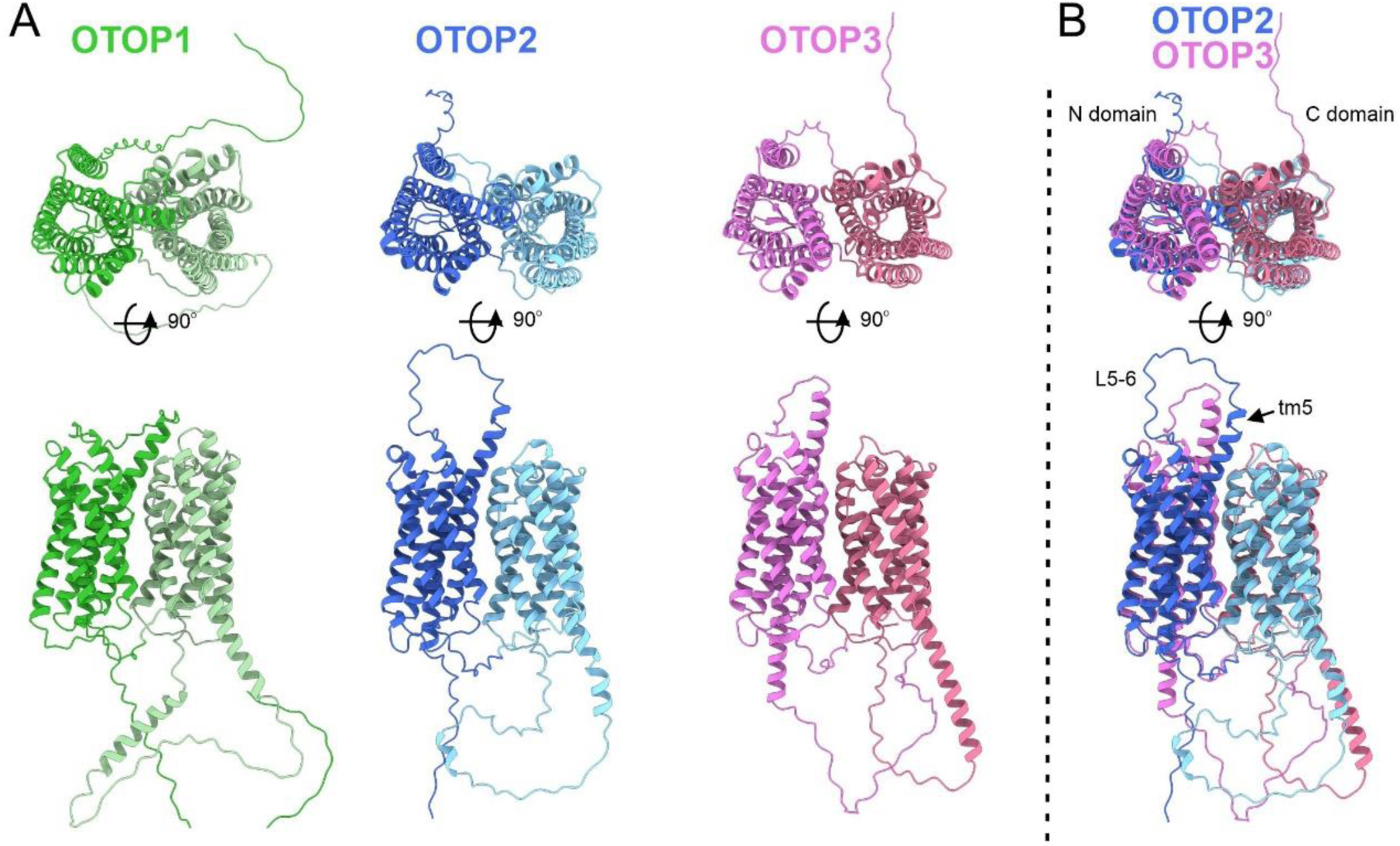
Predicted structures of mOTOP1. mOTOP2 and mOTOP3. **A**. Top views (top) and side views (bottom) of AlphaFold predicted structural models of mOTOP1, mOTOP2, and mOTOP3. The N- and C-domain halves of mOTOP1, mOTOP2, and mOTOP3 are colored green and light green, blue and light blue, and magenta and hot pink respectively. **B**. A superimposed overlay of mOTOP2 and mOTOP3 highlights the different orientations of the transmembrane 5 helices and S5-6 linkers.

## Discussion

Unlike other ion channels where functional descriptions of ion channel properties largely preceded their molecular identification, for OTOP1 and related proteins, such descriptions were limited to a few studies of the OTOP1 channels in taste receptor cells (Bushman, Ye, and Liman 2015; Chang, Waters, and Liman 2010). Thus, while ion selectivity of vertebrate and invertebrate OTOP channels has been well established (Tu et al. 2018; Chen et al. 2019), descriptions of basic gating mechanisms have been limited. For example, previous studies showed that the channels are not voltage-sensitive (Tu et al. 2018), but whether they are ligand-gated was not known. Here, using patch-clamp recording and pH imaging, we provide the first direct evidence that the three murine OTOP channels are gated by extracellular protons in a subtype-specific manner (Figure 9). OTOP1 and OTOP3 are proton-activated proton channels, while OTOP2 is a spontaneously open and proton-inhibited proton channel. The varying pH sensitivity of the three vertebrate OTOP channels may allow them to subserve different functions in the cells within which they are expressed and is reminiscent of the varying temperatures sensitivities of thermo-TRP channels (Dhaka, Viswanath, and Patapoutian 2006).

**Figure 9.**
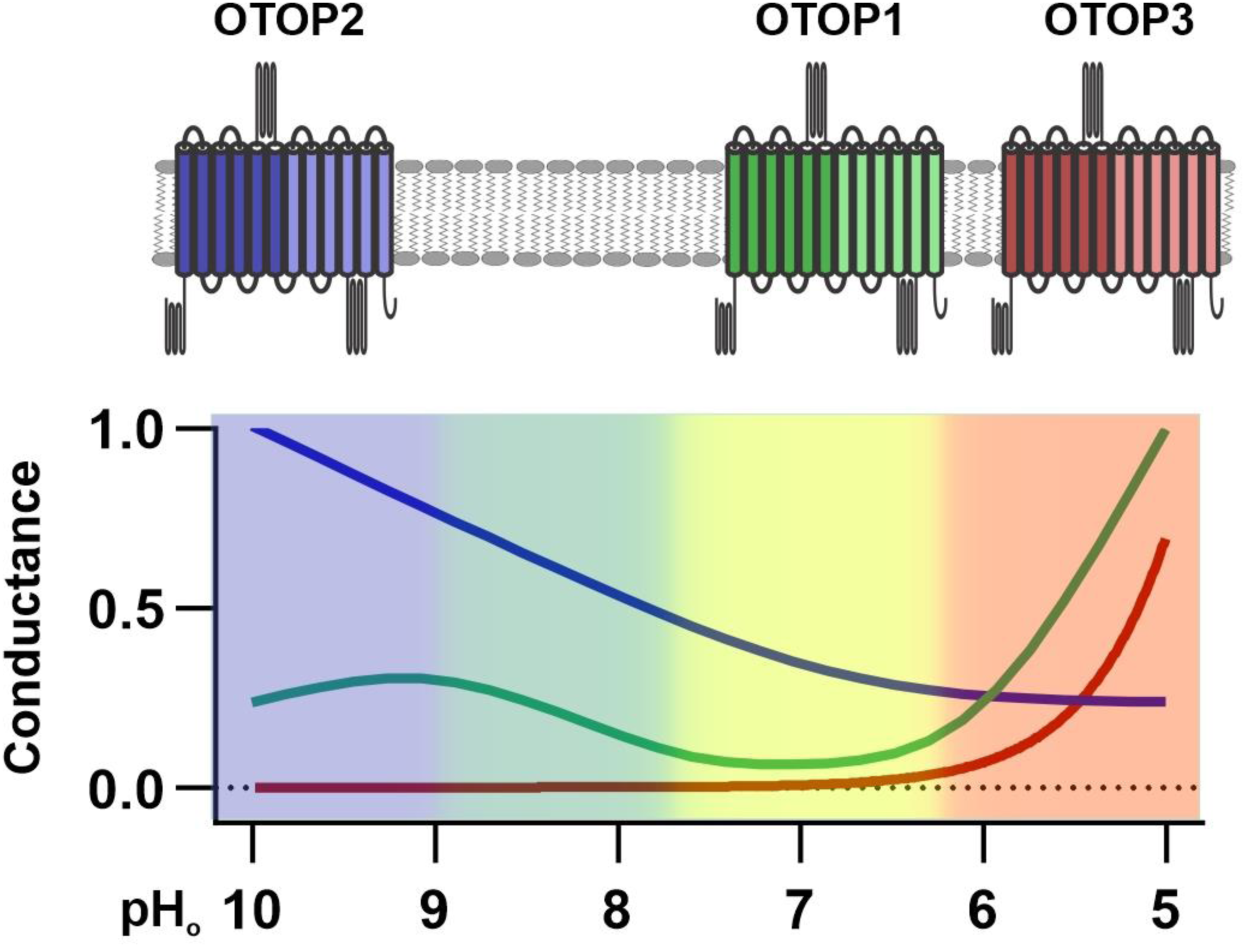
pH tuning of OTOP channels. Lowering extracellular pH increases the slope conductance of OTOP1 and OTOP3 channels while it lowers the slope conducance of OTOP2 channels. OTOP1 also has a peak of activity at mildly alkaline pH.

### Gating in response to changes in extracellular pH

The gating of ion channels by protons has been described previously. For example, the thermosensitive ion channel TRPV1, the acid-sensing ion channels (ASIC), and proton-activated chloride channels (PAC) are all activated by protons acting on extracellular sites (Waldmann et al. 1997; Jordt, Tominaga, and Julius 2000; Yang, Chen, et al. 2019; Ruan et al. 2020; Lambert and Oberwinkler 2005; Ullrich et al. 2019). Others are gated by protons acting on the intracellular side of the membrane (Cuello et al. 2010; Wang, Chang, and Liman 2010). Notably, the only other eukaryotic proton channel, the voltage-gated proton channel Hv1, shows a shift in voltage dependence as a function of pH (Ramsey et al. 2006; Decoursey 2012). For channels in which the proton is not the main permeant ion, the effect of pH on gating can be measured simply by comparing current magnitudes as extracellular pH is varied. In contrast, establishing an effect of pH (protons) on the gating of a proton channel is not so straightforward, and any analysis is further complicated by the exceedingly small conductance of proton channels at physiological or even acid pH (Decoursey 2012).

The magnitude of OTOP currents was previously shown to vary as a function of pHo (Tu et al. 2018). We confirmed the effects of pHo and extended the pH range of test solutions, allowing us to observe that OTOP2 and, to a lesser extent, OTOP1 channels can carry outward proton currents in response to extracellular alkalization. The differences in currents suggest underlying differences in gating. To provide evidence for differential gating, we devised a strategy that avoids confounds introduced by the changing driving force for proton movement inherent in these experiments. Taking advantage of differences in the three mammalian OTOP channels, we provide three independent pieces of evidence that extracellular protons gate the OTOP channels. First, we found that OTOP1 and OTOP3 currents activated slowly in response to acid stimuli, with rates that increased as the extracellular pH was lowered. These results can only be explained if lowering the pH opens the channels. Second, the slope conductance of the OTOP channels, which is independent of driving force, varied as a function of pHo for all three channels. Consistent with the kinetic data, the slope conductance of both OTOP1 and OTOP3 currents increased dramatically upon lowering the pH below 6.0 (OTOP1) and 5.5 (OTOP3), while the slope conductance of OTOP2 increased with alkalization above pH 8.0. Finally, these results are consistent with results from pH imaging experiments where proton efflux in response to extracellular alkalinization was only observed for OTOP1 and OTOP2 and not for OTOP3-expressing cells. Together these data allow us to conclude that the three murine OTOP channels are gated, differentially, by extracellular protons.

### Structure of OTOP channels and role of extracellular linkers

The recent cryo-EM structures of OTOP1 and OTOP3 channels have revealed that the channels adopt a novel fold, with twelve transmembrane alpha-helices organized into two structurally homologous six-helix domains (N and C domains) (Saotome et al. 2019; Chen et al. 2019). Thus, as a dimer, the channels adopt a pseudo-tetrameric structure. However, in contrast to most ion channels where the subunits come together to form a shared pore, the central cavity of OTOP channels is filled with cholesterol-like molecules and therefore cannot mediate ion conduction (Saotome et al. 2019). Three putative pathways for protons have been identified in the structures: one each in the N and C domains and another at the interface of N and C domains (intrasubunit interface). Mutations in each putative permeation pathway that cause loss of function have been identified, including mutations of conserved residues (E267 and H574 in ZfOTOP1) at the intrasubunit interface that are required for function, but not subunit assembly or trafficking (Saotome et al. 2019). Whether these residues are mediate gating or permeation of the channels is still not known, and it is possible they participate in both as is the case for Hv1 (Tombola, Ulbrich, and Isacoff 2008; Musset et al. 2011). Moreover, with three possible routes for proton permeation, there may be multiple pores that open under different conditions. For example, our observation that OTOP1 channels show a biphasic response to extracellular pH could be explained by two pores that open over different pH ranges.

Our data also provide a framework for understanding the structural basis for the gating of OTOP channels by protons. Swapping extracellular loops of OTOP2 and OTOP3 selectively changed gating of the channels in a predicable manner. Sapping loops in the N domain of the channels selectively affected the outward currents elicited in response to alkalinization, such that OTOP2/OTOP3 (L5-6) channels carried less outward current, while OTOP3/OTOP2(L3-4) channels carried more outward current that the respective wildtype channels. This suggests that the N domain either contains the pore that is active at alkaline pH or contains the alkaline pH sensor that opens a common pore (for example at the intrasubunit interface). In contrast, swapping loops within the C domain did not affect the gating of OTOP2 but changed the gating of OTOP3 channels such that they opened faster and at higher pH (like OTOP2). This suggests that the extracellular loops in the C domain of OTOP3 inhibit activation of the channels; moreover, the observation that nearly all swaps led to an increase in the rate of current activation suggests that the slow gating kinetics characteristic of OTOP3 is the result of multiple domains acting in concert. Although the structure of OTOP2 has not been resolved experimentally, structural predictions from Alphafold (Varadi et al. 2022; Jumper et al. 2021) point to intriguing differences in the position of S5 and the S5-6 linker between OTOP2 and OTOP3 that may underlie differences in gating.

Together our description of the differential gating of the three mammalian OTOP channels by protons provides the basis for a deeper understanding of this new ion channel family. Further understanding the mechanism by which protons gate OTOP channels will require a combination of structure-guided mutagenesis and the resolution of the structure of OTOP channels in different states (Ruan et al. 2020; Deng et al. 2021). Our description of the differential pH dependence of the three vertebrate OTOP channels and the unique features of OTOP2 channels will lead to a better understanding of how OTOP channels contribute to a wide range of physiological processes, from gravity senses and taste to digestion, both in health and disease.

## Materials and Methods

### Clones, cell lines, and transfection

Mouse Otop1, Otop2, and Otop3 cDNAs were cloned into the pcDNA3.1 vector as previously described (Tu et al. 2018). For experiments in Figure 6 and 7, both mOtop2 and mOtop3 cDNAs were cloned into the pcDNA3.1 vector with an N-terminal fusion tag consisting of an octahistidine tag followed by eGFP, a Gly-Thr-Gly-Thr linker, and 3C protease cleavage site (LEVLFQGP) as previously described (Saotome et al. 2019). Chimeras were generated using In-Fusion Cloning (Takara Bio) and were confirmed by Sanger sequencing (Genewiz).

HEK293 cells were purchased from ATCC (CRL-1573). PAC-KO cells were a kind gift from Dr. Zhaozhu Qiu (Yang, Chen, et al. 2019). The cells were cultured in a humidified incubator at 37C in 5% CO2 and 95% O2 using a high glucose DMEM (ThermoFisher) containing 10% fetal bovine serum (Life Technology) and 1% Penicillin-streptomycin antibiotics. Cells were passaged every 3 – 4 days.

Cells used for patch-clamp recordings were transfected in 35mm Petri dishes, with ∼ 600ng DNA and 2 µL TransIT-LT1 transfection reagents (Mirus Bio Corporation) following the manufacturer’s protocol. OTOP channels were co-transfected with GFP or pHluorin at a ratio of 5:1. The cells were lifted using Trypsin-EDTA 24 h after transfection, plated onto a coverslip, and used within 3-4 hrs for patch-clamp recordings.

### Patch-clamp electrophysiology

Whole-cell patch-clamp recording was performed as previously described (Tu et al. 2018). Briefly, recordings were obtained with an Axonpatch 200B amplifier, digitized with a Digidata 1322a 16-bit data acquisition system, acquired with pClamp 8.2, and analyzed with Clampfit 9 (Molecular Devices). Patch pipettes with a resistance of 2 – 4 MΩ were fabricated from borosilicate glass (Sutter instrument) and fire polished. Only cells with stable giga-ohm seals were used for data collection and subsequent analysis. Records were sampled at 5 kHz and filtered at 1 kHz. Millisecond solution exchange was achieved with a fast-step perfusion system (Warner instrument, SF-77B) custom modified to hold seven microcapillary tubes in a linear array.

The holding potential was -80 mV unless otherwise indicated. For experiments in Figure 5, the voltage was held at -80 mV and ramped to +80 m at 1 V/s, once per second. The slope of the first ramp after the peak current in response to acids was measured to determine the conductance.

### Patch-clamp electrophysiology solutions

Tyrode’s solution contained 145 mM NaCl, 5 mM KCl, 1 mM MgCl2, 2 mM CaCl2, 20mM dextrose, 10mM HEPES (pH adjusted to 7.4 with NaOH). Standard pipette solution contained 120 mM Cs-aspartate, 15 mM CsCl, 2 mM Mg-ATP, 5 mM EGTA, 2.4 mM CaCl2 (100nM free Ca2+), and 10 mM HEPES (pH adjusted to 7.3 with CsOH; 290 mosm). Standard Na+ free extracellular solutions contained 160 mM NMDG-Cl, 2 mM CaCl2, and 10 mM buffer based on pH (CHES for pH 10 – 9, HEPES for pH 8 – 7.4, PIPES for pH 7, Bis-tris for pH 6.5, MES for pH 6 - 5.5 and HomoPIPES for pH 5 - 4.5).

For sodium solution in Figure 2A, 160 mM NMDG-Cl was replaced by an equimolar concentration of NaCl. For the low chloride solution in Figure 2B, 120 mM HCl was replaced by methane sulfonic acid (CH3SO3H). In these experiments, 200 µM Amiloride was added to block endogenous ASIC channels.

### pH imaging for transfected HEK-293 cells

HEK-293 cells were cultured in 35 mm Petri dishes. OTOP channels and the pH-sensitive indicator pHluorin were co-transfected into the cells. After 24 hours, the cells were lifted and plated on poly-D-lysine coated coverslips at room temperature. The cells were incubated in standard Tyrodes’ solutions before experiments, and the pHluorin fluorescence intensity in response to different solutions was measured. All stimulating solutions were modified from the standard Tyrodes’ solution, containing 10 mM of different buffer salt based on the pH (CHES for pH 8.5, MES for pH 6.0, Homo-PIPES for pH 5.0, and Acetic acid for pH 5.0 [HOAC] group). Excitation was at 488 nm, and emission was detected at 510 nm using a U-MNIBA2 GFP filter cube (Olympus). Images were acquired on a Hamamatsu digital CCD camera attached to an Olympus IX71 microscope using Simple PCI software. The fluorescence intensity of each cell was normalized to its baseline in Tyrodes’ solutions (F0) before the first test stimulus was given to the cells.

### Quantification, statistical analysis, and generation of AlphaFold models

All data are presented as mean ± SEM if not otherwise noted. Statistical analysis was performed using Graphpad Prism 8 or 9 (Graphpad Software Inc). The sample sizes of 3-10 independent recordings from individual cells per data point are similar to those in the literature for similar studies. All data are biological replicates. In some cases where technical replicates were performed (e.g. re-running the test protocol a second time on the same cell), we typically used the first series, unless there was a loss of seal resistance and recovery, in which case we used the second replicate. Selection, in this case, was blind to the result/outcome. Representative electrophysiology traces shown in the figures were acquired with pClamp, and in some cases, the data was decimated by 10-fold before exporting into graphic programs, Origin (Microcal) and Coreldraw (Corel).

The predicted structural models of mOTOP1, mOTOP2, and mOTOP3 were downloaded from the AlphaFold Protein Structure Database https://alphafold.ebi.ac.uk (Varadi et al. 2022; Jumper et al. 2021). Figures of the models were generated using UCSF ChimeraX https://www.rbvi.ucsf.edu/chimerax/ (Goddard et al. 2018).

## Acknowledgments

We thank Jackson Walker and Anne Tran for expert technical support and all members of the Liman and Ward labs for helpful discussions. We also thank M. Goldschen-ohm for critical reading of the manuscript, and Z. Qui for generously sharing cell lines. Research reported in this publication was supported by the National Institute of General Medical Sciences of the NIH under award number R01GM131234 and from the National Institute for Deafness and Communication Disorders under award number R01DC013741 to E.R.L.

## Competing interests

The authors declare no competing interests

---

**Supplementary Figure 1 (related to Fig 4).**
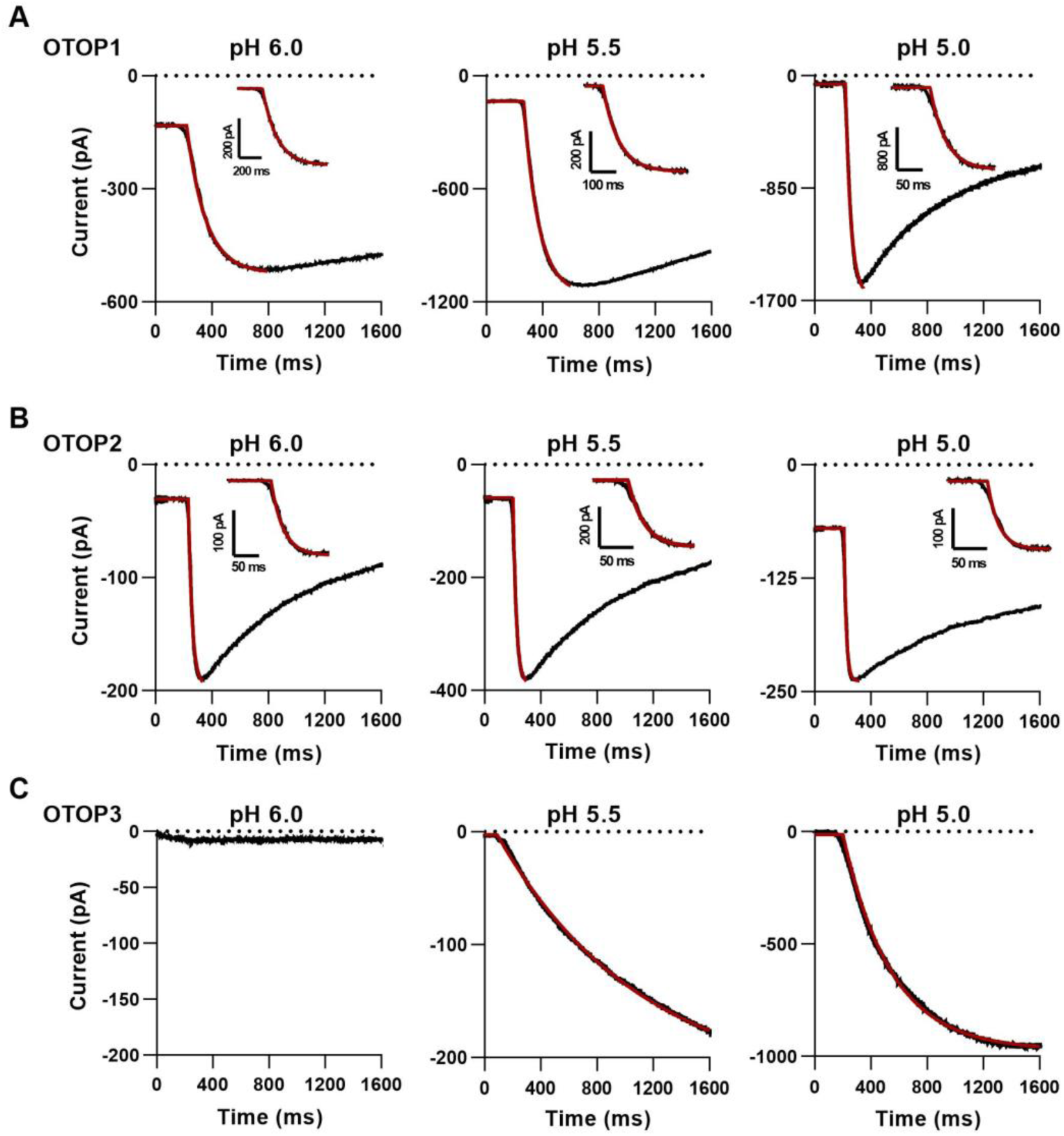
Fit to activation kinetics of OTOP currents. **A-C** Representative traces showing the activation of OTOP1 (A), OTOP2 (B), and OTOP3 (C) currents in response to solutions with extracellular pH titrated to 6.0 (left panel), 5.5 (middle panel), and 5.0 (right panel). The activation of the currents was fit by a single exponential curve (red part of the trace). Inserted shows a zoom-in view of the current activation. The initial few milliseconds where data deviated from an exponential were not included in the curve fitting as they reflect the non-instantaneous solution exchange. No measurable current was evoked in OTOP3 in response to pH 6.0.

**Supplementary Figure 2.**
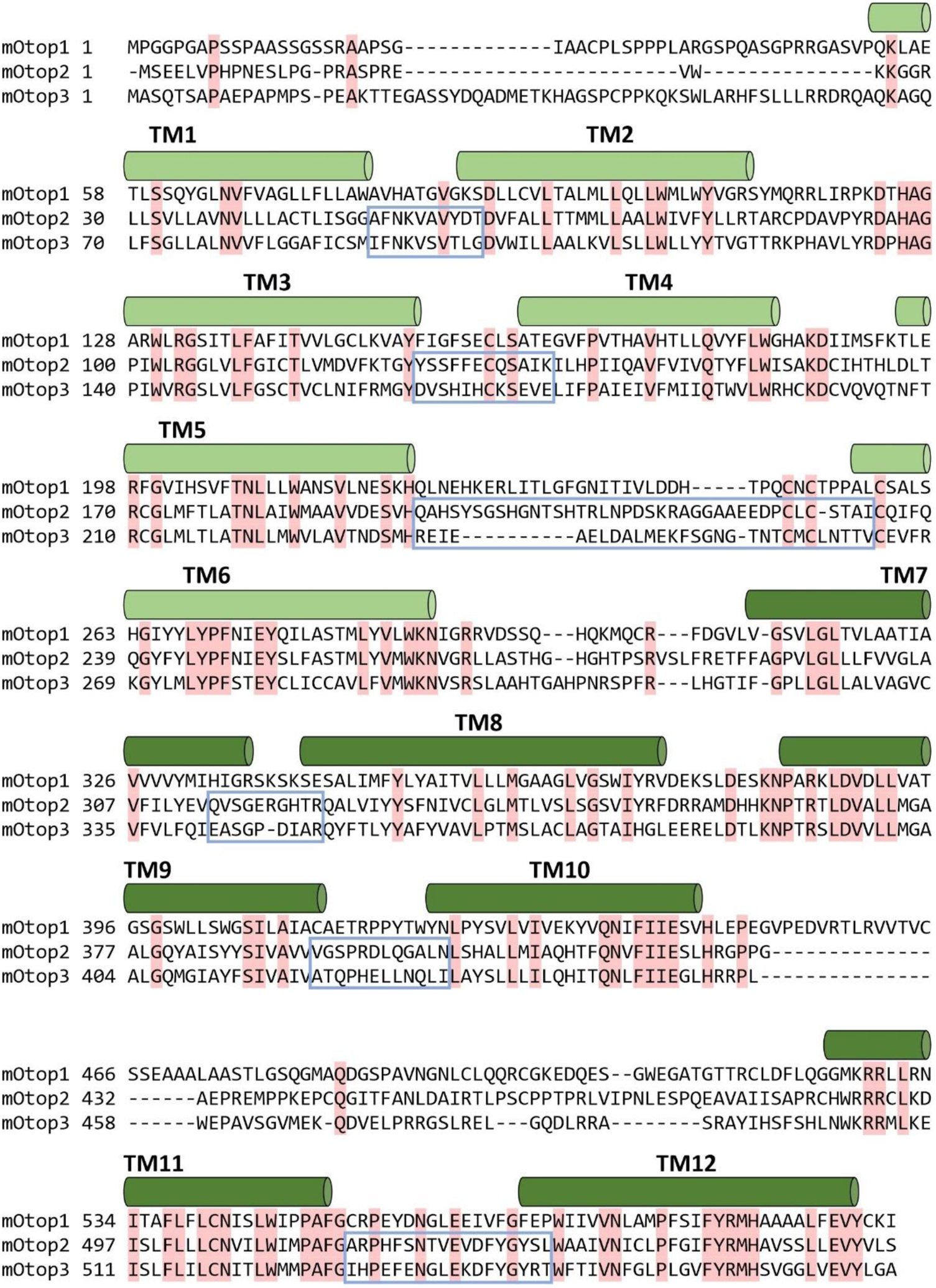
Topology and sequence alignment of OTOP channels. Mouse OTOP1, 2, and 3 channels are aligned. The transmembrane helices are depicted in cylinders above the sequence. Residuals that are conserved in all three channels are labeled with pink shade. External loops that were swapped in OTOP2 and OTOP3 are labeled in blue boxes.

**Supplementary Table 1.** Statistical tests comparing chimeric channels with wildtype channels with Mann-Whitney test. P values and sample size are indicated.

|  | OTOP2 |  | OTOP3 |  |
| --- | --- | --- | --- | --- |
|  | pH 10 | pH 6 | pH 10 | pH 6 |
| WT | n=4 | n=4 | n=5 | n=5 |
| L1-2 | 0.063492<br>n=5 | 0.730159<br>n=5 | NA | NA |
| L3-4 | NA | NA | **0.015873**<br>n=4 | **0.035714**<br>n=3 |
| L5-6 | **0.028571**<br>n=4 | **0.028571**<br>n=4 | 0.063492<br>n=4 | **0.015873**<br>n=4 |
| L7-8 | 0.685714<br>n=4 | 0.999999<br>n=4 | 0.690476<br>n=5 | **0.015873**<br>n=5 |
| L9-10 | NA | NA | 0.84127<br>n=5 | **0.007937**<br>n=5 |
| L11-12 | 0.171429<br>n=6 | 0.761905<br>n=6 | 0.555556<br>n=4 | **0.004329**<br>n=6 |

\* NA indicates no measurable currents.
\*\*The numbers are labeled red if $P < 0.05$ .

## Notes

### Competing Interest Statement

The authors have declared no competing interest.

## References

Bushman, J. D., W. Ye, and E. R. Liman. 2015. ‘A proton current associated with sour taste: distribution and functional properties’, Faseb J, 29: 3014–26.

Chang, R. B., H. Waters, and E. R. Liman. 2010. ‘A proton current drives action potentials in genetically identified sour taste cells’, Proc Natl Acad Sci U S A, 107: 22320–5.

Chang, W. W., A. S. Matt, M. Schewe, M. Musinszki, S. Grussel, J. Brandenburg, D. Garfield, M. Bleich, T. Baukrowitz, and M. Y. Hu. 2021. ‘An otopetrin family proton channel promotes cellular acid efflux critical for biomineralization in a marine calcifier’, Proc Natl Acad Sci U S A, 118.

Chen, Q., W. Zeng, J. She, X. C. Bai, and Y. Jiang. 2019. ‘Structural and functional characterization of an otopetrin family proton channel’, Elife, 8.

Cuello, L. G., D. M. Cortes, V. Jogini, A. Sompornpisut, and E. Perozo. 2010. ‘A molecular mechanism for proton-dependent gating in KcsA’, FEBS Lett, 584: 1126–32.

Decoursey, T. E. 2003. ‘Voltage-gated proton channels and other proton transfer pathways’, Physiol Rev, 83: 475–579.

Decoursey, T. E.. 2012. ‘Voltage-gated proton channels’, Compr Physiol, 2: 1355–85.

Deng, Z., Y. Zhao, J. Feng, J. Zhang, H. Zhao, M. J. Rau, J. A. J. Fitzpatrick, H. Hu, and P. Yuan. 2021. ‘Cryo-EM structure of a proton-activated chloride channel TMEM206’, Sci Adv, 7.

Dhaka, A., V. Viswanath, and A. Patapoutian. 2006. ‘Trp ion channels and temperature sensation’, Annu Rev Neurosci, 29: 135–61.

Ganguly, A., A. Chandel, H. Turner, S. Wang, E. R. Liman, and C. Montell. 2021. ‘Requirement for an Otopetrin-like protein for acid taste in Drosophila’, Proc Natl Acad Sci U S A, 118.

Goddard, T. D., C. C. Huang, E. C. Meng, E. F. Pettersen, G. S. Couch, J. H. Morris, and T. E. Ferrin. 2018. ‘UCSF ChimeraX: Meeting modern challenges in visualization and analysis’, Protein Sci, 27: 14–25.

Hille, B. 2001. Ionic channels of excitable membranes (Sinauer Associates Inc.: Sunderland, MA).

Hughes, I., B. Blasiole, D. Huss, M. E. Warchol, N. P. Rath, B. Hurle, E. Ignatova, J. D. Dickman, R. Thalmann, R. Levenson, and D. M. Ornitz. 2004. ‘Otopetrin 1 is required for otolith formation in the zebrafish Danio rerio’, Dev Biol, 276: 391–402.

Hurle, B., E. Ignatova, S. M. Massironi, T. Mashimo, X. Rios, I. Thalmann, R. Thalmann, and D. M. Ornitz. 2003. ‘Non-syndromic vestibular disorder with otoconial agenesis in tilted/mergulhador mice caused by mutations in otopetrin 1’, Hum Mol Genet, 12: 777–89.

Hurle, B., T. Marques-Bonet, F. Antonacci, I. Hughes, J. F. Ryan, Nisc Comparative Sequencing Program, E. E. Eichler, D. M. Ornitz, and E. D. Green. 2011. ‘Lineage-specific evolution of the vertebrate Otopetrin gene family revealed by comparative genomic analyses’, BMC Evol Biol, 11: 23.

Jordt, S. E., M. Tominaga, and D. Julius. 2000. ‘Acid potentiation of the capsaicin receptor determined by a key extracellular site’, Proc Natl Acad Sci U S A, 97: 8134–9.

Jumper, J., R. Evans, A. Pritzel, T. Green, M. Figurnov, O. Ronneberger, K. Tunyasuvunakool, R. Bates, A. Zidek, A. Potapenko, A. Bridgland, C. Meyer, S. A. A. Kohl, A. J. Ballard, A. Cowie, B. Romera-Paredes, S. Nikolov, R. Jain, J. Adler, T. Back, S. Petersen, D. Reiman, E. Clancy, M. Zielinski, M. Steinegger, M. Pacholska, T. Berghammer, S. Bodenstein, D. Silver, O. Vinyals, A. W. Senior, K. Kavukcuoglu, P. Kohli, and D. Hassabis. 2021. ‘Highly accurate protein structure prediction with AlphaFold’, Nature, 596: 583–89.

Lambert, S., and J. Oberwinkler. 2005. ‘Characterization of a proton-activated, outwardly rectifying anion channel’, J Physiol, 567: 191–213.

Liang, R., J. M. J. Swanson, J. J. Madsen, M. Hong, W. F. DeGrado, and G. A. Voth. 2016. ‘Acid activation mechanism of the influenza A M2 proton channel’, Proc Natl Acad Sci U S A, 113: E6955–E64.

Mi, T., J. O. Mack, C. M. Lee, and Y. V. Zhang. 2021. ‘Molecular and cellular basis of acid taste sensation in Drosophila’, Nat Commun, 12: 3730.

Miesenbock, G., D. A. De Angelis, and J. E. Rothman. 1998. ‘Visualizing secretion and synaptic transmission with pH-sensitive green fluorescent proteins’, Nature, 394: 192–5.

Morgan, D., M. Capasso, B. Musset, V. V. Cherny, E. Rios, M. J. Dyer, and T. E. DeCoursey. 2009. ‘Voltage-gated proton channels maintain pH in human neutrophils during phagocytosis’, Proc Natl Acad Sci U S A, 106: 18022–7.

Musset, B., S. M. Smith, S. Rajan, D. Morgan, V. V. Cherny, and T. E. Decoursey. 2011. ‘Aspartate 112 is the selectivity filter of the human voltage-gated proton channel’, Nature, 480: 273–7.

Parikh, K., A. Antanaviciute, D. Fawkner-Corbett, M. Jagielowicz, A. Aulicino, C. Lagerholm, S. Davis, J. Kinchen, H. H. Chen, N. K. Alham, N. Ashley, E. Johnson, P. Hublitz, L. Bao, J. Lukomska, R. S. Andev, E. Bjorklund, B. M. Kessler, R. Fischer, R. Goldin, H. Koohy, and A. Simmons. 2019. ‘Colonic epithelial cell diversity in health and inflammatory bowel disease’, Nature, 567: 49–55.

Pinto, L. H., L. J. Holsinger, and R. A. Lamb. 1992. ‘Influenza virus M2 protein has ion channel activity’, Cell, 69: 517–28.

Qu, H., Y. Su, L. Yu, H. Zhao, and C. Xin. 2019. ‘Wild-type p53 regulates OTOP2 transcription through DNA loop alteration of the promoter in colorectal cancer’, FEBS Open Bio, 9: 26–34.

Ramsey, I. S., M. M. Moran, J. A. Chong, and D. E. Clapham. 2006. ‘A voltage-gated proton-selective channel lacking the pore domain’, Nature, 440: 1213–6.

Ruan, Z., J. Osei-Owusu, J. Du, Z. Qiu, and W. Lu. 2020. ‘Structures and pH-sensing mechanism of the proton-activated chloride channel’, Nature, 588: 350–54.

Saotome, K., B. Teng, C. C. A. Tsui, W. H. Lee, Y. H. Tu, J. P. Kaplan, M. S. P. Sansom, E. R. Liman, and A. B. Ward. 2019. ‘Structures of the otopetrin proton channels Otop1 and Otop3’, Nat Struct Mol Biol, 26: 518–25.

Sasaki, M., M. Takagi, and Y. Okamura. 2006. ‘A voltage sensor-domain protein is a voltage-gated proton channel’, Science, 312: 589–92.

Teng, B., C. E. Wilson, Y. H. Tu, N. R. Joshi, S. C. Kinnamon, and E. R. Liman. 2019. ‘Cellular and Neural Responses to Sour Stimuli Require the Proton Channel Otop1’, Curr Biol, 29: 3647–56 e5.

Tombola, F., M. H. Ulbrich, and E. Y. Isacoff. 2008. ‘The voltage-gated proton channel Hv1 has two pores, each controlled by one voltage sensor’, Neuron, 58: 546–56.

Tu, Y. H., A. J. Cooper, B. Teng, R. B. Chang, D. J. Artiga, H. N. Turner, E. M. Mulhall, W. Ye, A. D. Smith, and E. R. Liman. 2018. ‘An evolutionarily conserved gene family encodes proton-selective ion channels’, Science, 359: 1047–50.

Ullrich, F., S. Blin, K. Lazarow, T. Daubitz, J. P. von Kries, and T. J. Jentsch. 2019. ‘Identification of TMEM206 proteins as pore of PAORAC/ASOR acid-sensitive chloride channels’, Elife, 8.

Varadi, M., S. Anyango, M. Deshpande, S. Nair, C. Natassia, G. Yordanova, D. Yuan, O. Stroe, G. Wood, A. Laydon, A. Zidek, T. Green, K. Tunyasuvunakool, S. Petersen, J. Jumper, E. Clancy, R. Green, A. Vora, M. Lutfi, M. Figurnov, A. Cowie, N. Hobbs, P. Kohli, G. Kleywegt, E. Birney, D. Hassabis, and S. Velankar. 2022. ‘AlphaFold Protein Structure Database: massively expanding the structural coverage of protein-sequence space with high-accuracy models’, Nucleic Acids Res, 50: D439–D44.

Waldmann, R., G. Champigny, F. Bassilana, C. Heurteaux, and M. Lazdunski. 1997. ‘A proton-gated cation channel involved in acid-sensing’, Nature, 386: 173–7.

Wang, Y. Y., R. B. Chang, S. D. Allgood, W. L. Silver, and E. R. Liman. 2011. ‘A TRPA1-dependent mechanism for the pungent sensation of weak acids’, J Gen Physiol, 137: 493–505.

Wang, Y. Y., R. B. Chang, and E. R. Liman. 2010. ‘TRPA1 is a component of the nociceptive response to CO2’, J Neurosci, 30: 12958–63.

Yang, H., H. Liu, H. C. Lin, D. Gan, W. Jin, C. Cui, Y. Yan, Y. Qian, C. Han, and Z. Wang. 2019. ‘Association of a novel seven-gene expression signature with the disease prognosis in colon cancer patients’, Aging (Albany NY), 11: 8710–27.

Yang, J., J. Chen, M. Del Carmen Vitery, J. Osei-Owusu, J. Chu, H. Yu, S. Sun, and Z. Qiu. ‘PAC, an evolutionarily conserved membrane protein, is a proton-activated chloride channel’, Science, 364: 395–99.

Zhang, J., H. Jin, W. Zhang, C. Ding, S. O’Keeffe, M. Ye, and C. S. Zuker. 2019. ‘Sour Sensing from the Tongue to the Brain’, Cell, 179: 392–402 e15.

